# Chromatin-interacting transposon RNAs linking to the core trans-inhibition circuitry for embryonic stem cell identity

**DOI:** 10.1101/2021.04.28.441894

**Authors:** Wenqiu Xu, Likun Ren, Caihong Zheng, Jun Cai

## Abstract

Transposable DNA sequences constitute more than half of the human and mouse genomes. A large number of non-coding RNAs, named as transposon RNAs, are derived from these transposable elements. The cis-regulatory function of transposable DNA elements, such as LINE-1 and Alu has been largely explored. But the biological roles of transposon RNAs aren’t well understood. Here, investigations of RNA-chromatin interactions provide us with comprehensive evidence that specific families of transposon RNAs play roles in trans-regulation linking to the core inhibition circuitry for embryonic stem cell identity. Alternative modes of the RNA-DNA hybrid duplex and protein-recruited RNA scaffold are required for the regulatory activities of transposon RNAs. LINE-1 RNAs co-locating with KAP1 form a negative feedback loop stabilizing the transcription of LINE-1 DNA elements via RNA-DNA hybrids. In another way LINE-1 RNAs, together with the reprogramming three factors and Polycomb repressive complexes, participate in the inhibition on dozens of differentiation-relative genes.

## Introduction

Through independent replication of their sequences, transposable elements (TEs) become a group of the most abundant genetic materials within the genome. They account for approximately 46% of the human genome and 37.5% of mouse genome, respectively (Mouse Genome Sequencing et al. 2002). Such repetitive sequences were initially considered as ‘selfish’ and ‘junk’ genetic elements, whose expression or mobility could be silenced in cells by some epigenetic mechanisms such as DNA methylation (Gaudet et al. 2004; Arand et al. 2012) and histone modification (Bulut-Karslioglu et al. 2014). Increasingly, researchers have realized the importance of these transposable DNA sequences as a potent source of cis-regulatory DNA motifs to recruit specific regulatory factors, regulate the functional activities of coding genes (Bourque et al. 2008; Kunarso et al. 2010; Testori et al. 2012; Sundaram et al. 2014) and establish the 3D chromatin structure (Lu et al. 2021). For example, Alu sequences, acting as transcription factor binding sites or enhancer elements, are cis-regulating diverse RNA processes, such as RNA transcription, RNA editing, and exonization (Mandal et al. 2013). Stage-specific endogenous retroviruses (ERVs) as DNA regulator elements activate gene expression during pre-implantation embryo development (Fu et al. 2019). Further studies on long non-coding RNAs (lncRNAs) gained a deeper understanding of TEs. The TEs are not only major signals essential for the origin and diversification of lncRNAs, but also contribute to regulation roles of many lncRNAs (Kelley and Rinn 2012). A growing number of TE insertion regulatory domains in lncRNAs, which act as RNA-, DNA-, and protein-binding domains, have been experimentally defined (Johnson and Guigo 2014). But, except for the individual experimental evidence that Alu elements in *ANRIL* noncoding RNA modulate cell function through trans-regulation of gene networks (Holdt et al. 2013), the role of TEs was still focused on their cis-acting functions on regulating the transcription initiation, splicing, or polyadenylation.

The expressed transposon RNAs were thought to be detrimental and cause mutations to cells (Malki et al. 2014; Burns 2017). When a series of small- and large-scale experiments captured the engagement of TEs in cis-regulatory activities, we seem to have lost sight of the involvement of trans-regulatory activities for TE-derived transposon RNAs. Fortunately, the current development of advanced large-scale experimental biotechnologies, such as MARGI technology (Sridhar et al. 2017), GRID-seq (Li et al. 2017), ChAR-seq (Bell et al. 2018), and RADICL-seq (Bonetti et al. 2020), enables the whole-transcriptome profiling of RNA-chromatin interactions. These high-throughput experimental data provide a special opportunity for us to explore the interactions between transposon-derived RNAs and chromatin for the understanding of the potential trans-regulatory function of these transposon RNAs. Researchers initiated to utilize the RNA-chromatin interaction data for the discovery that transposon RNAs modulate the process of heterochromatin organization (Hao et al. 2020; Chen et al. 2021; Liu et al. 2021). In this study, we performed a high-throughput proximity MARGI experiment in E14 cells and interpreted the roles of transposon RNAs from the perspective of RNA trans-regulation on gene transcription in embryonic stem cells (ESCs). We concern about the answers to a series of questions: which subfamilies of transposon RNAs prefer to target the genomic regions? what kind of binding modes drive the interactions between transposon RNAs and chromatins? and how do the transposon RNAs trans-regulate the gene expression of the core regulatory circuitry for ESC identity?

## Results

### Preferred families of transposon RNAs interacting with chromatins

We performed a high-throughput MARGI experiment in mouse E14 cells and integrated the public MARGI data in human H9 and HEK293 cells for analysis. Effective and reasonable criteria were utilized to define the distal transposon RNA-chromatin interaction distinct from the local interaction between RNA and its coding DNA sequence (Sridhar et al. 2017) (See Methods). The data analysis results confirm the high reproducibility and reliability of the distal interactions defined by reduplicate MARGI experiments in E14 cells (Figure 1A). The distal interactions between transposon RNAs and chromatins in H9, HEK293, and E14 are explored, respectively (Figure 1B, Figure S1A). On averaged, 91.28% distal transposon RNA-chromatin interactions are inter-chromosomal. Various TEs such as LINE, LTR, and SINE, as potential trans-regulating RNAs, are involved in the interactions with chromosomal DNAs (Figure 1C, Figure S1B). For the conserved LINE and LTR in mammal species, the LINE-1 family in LINE and ERV family in LTR prefer to interact with chromatins in both human and mouse (Figure 1C, Figure S1B).

**Figure 1.**
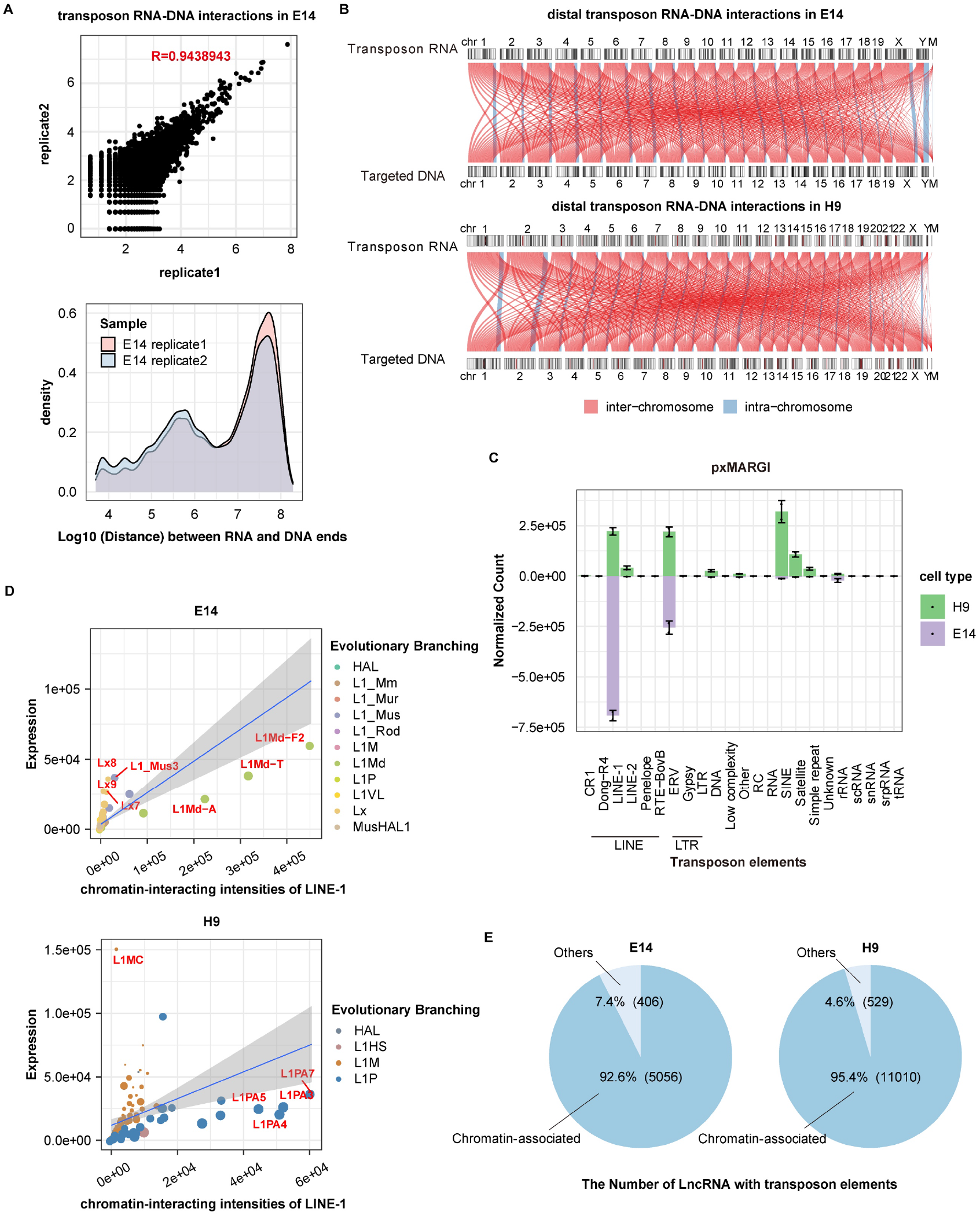
The characteristics of chromatin-associated transposon RNAs. (A) The repeatability of distal transposon RNA-chromatin interactions detected in E14 MARGI replicates (upper subplot); And the distribution of the coordinate distance between RNA and paired DNA in each intra-chromosome distal transposon RNA-chromatin interaction (lower subplot). In the upper subplot, the count of paired RNA-DNA reads supporting distal transposon RNA-chromatin interactions was calculated in each bin with 1kb length along the mouse genome. Each scatter point shows the two counts in the same bin for the two replicated experiments. Pearson correlation coefficient was calculated. (B) The connection map of distal transposon RNA-chromatin interactions across chromosomes in E14 and H9 cells. Each line represents a group of intra-chromosome (in blue color) or inter-chromosome (in red color) transposon RNA-chromatin interaction events between a pair of chromosomes. The thickness of the line indicates the number of interactions. (C) Preferred classes and families of chromatin-associated transposon RNAs across species. The bars show the mapped read counts of various transposon RNA families in distal interactions. Error bars represent standard error of replications. (D) The enrichment of LINE-1 RNA subfamilies involved in distal transposon RNA-chromatin interactions in E14 and H9 cells. The expression dosages in y-axis were used to normalize the chromatin-interacting intensities of LINE-1 RNA subfamilies. The preferred LINE-1 RNA subfamilies, being the outliers to the linear fitting curve of all the scatter points, are marked in red label. The different size and color of scatter points represent the evolutionary age and the specific evolutionary branches for each LINE-1 subfamily, respectively. (E) The proportion of lncRNAs with chromatin-interacting TE domains. The lncRNAs with TE insertion domains were annotated in Ensembl and GENCODE.

Considering the fact that the expressed copies of various transposon RNAs in different species or cell types are distinct, we normalized the chromatin-interacting intensities of transposon RNAs in subfamilies to ensure removal of the effect of RNA copies. In Figure 1D, the relative distance of each scatter point, to the linear fitting curve of all the scatter points describes the normalized chromatin-interacting intensity for one LINE-1 subfamily, such as HAL1, L1PA8, and L1Md-A. While the scatter point moves towards the lower right orientation, stronger chromatin-associated interaction occurs in the corresponding subfamily of L1PA3, L1PA4, L1PA5 or L1PA7 in human or L1Md in mouse. When we incorporate the evolutionary ages of TEs (Giordano et al. 2007) into the Figure 1D, it is found that the LINE-1 subfamilies interacting with chromatins in human and mouse cells are concentrated in the younger branches of L1PA and L1Md (Figure 1D, Figure S1C).

While a growing number of TE insertions in lncRNAs have been experimentally defined as regulatory domains (Johnson and Guigo 2014), the answer is yet unknown how many such TE domains in expressed lncRNAs indeed act as DNA-binding regulatory domains. About 80% of lncRNAs have TE insertion domains. We find that, in human H9 and HEK293 cells, 95% of TE insertion domains in lncRNAs execute the activity of DNA binding on the basis of sequencing evidence of interacting TE-chromatin pairs. But these TE regulatory domains account for less than 8% of the total chromatin-interacting TEs (Figure 1E, Figure S1D). In mouse E14 cells, about 92% of TE insertion domains in lncRNAs have the role of DNA binding, accounting for only 4% of the total chromatin-interacting TEs (Figure 1E). These results suggest that trans-regulation of transposon RNAs on chromosomal DNAs far outweighs the effect of TE regulatory domains in lncRNAs.

### Optional binding modes between transposon RNAs and chromatin regions

We extracted the nucleotide sequence contexts of the distal interactions between transposon RNAs and chromatins in H9, HEK293, and E14. RNA-DNA Watson-Crick complementary duplex and triple-helix structures formed by the sequence context of the paired transposon RNA and its targeting DNA region were predicted. DNA sequences outside the transposon RNA targeting regions were randomly sampled to compose the pseudo transposon RNA-DNA pairs as control. And the distal interaction pairs between non-transposon RNAs and chromatins were selected as another control group. The distribution of minimal free energy defining the binding strength of RNA-DNA hybrid duplexes in the sequence contexts of the paired ERV or LINE-1 RNA and its targeting DNA region has the unique characteristics of long tail and bimodal (Figure 2A). Quite numbers of interactions between ERV or LINE-1 RNAs and chromatins are significantly different from the two background control groups in the distributions with a false positive rate of 5% as the cutoff value. These ERV or LINE-1 RNA-chromatin interactions form RNA-DNA hybrids with ultra-low minimal free energy less than −60 kcal/mol, the binding strength of which is competitive with that of the RNA-DNA hybrids formed by pairwise perfect-matched sequences (Figure 2A). The existence of homolog repetitive elements of ERV or LINE-1 in the targeted DNA regions contributes to the formation of stable RNA-DNA hybrid duplexes in ERV or LINE-1 RNA-chromatin interactions. In total, stable Watson-Crick base pairing RNA-DNA hybrid duplexes mediate 30% ERV RNA-chromatin interactions and 40% LINE-1 RNA-chromatin interactions, respectively (Figure 2B). Few triplex-helix structures, on the contrary, occur in the sequence contexts of transposon RNA-chromatin interactions.

**Figure 2.**
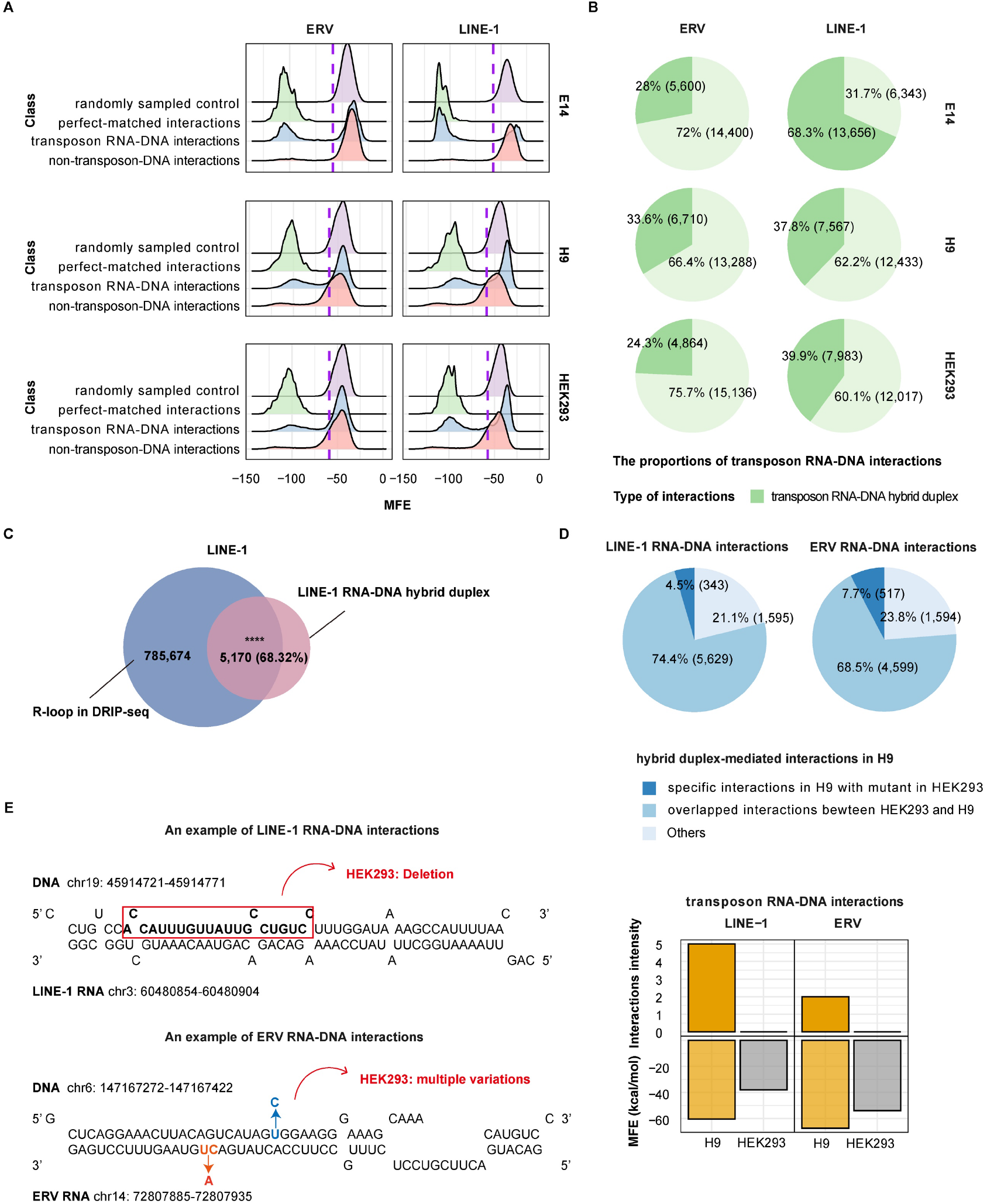
The interacting mode of RNA-DNA hybrids between transposon RNAs and chromatin regions. (A) The bimodal density distribution of minimal free energy. The minimal free energy defines the binding strength of a RNA-DNA hybrid duplex in the sequence context of each ERV or LINE-1 RNA and its pairwise targeting DNA region in E14, H9 and HEK293 cells. DNA sequences outside the transposon RNA targeting regions were randomly sampled to compose the pseudo transposon RNA-DNA pairs as control. And the distal interaction pairs between non-transposon RNAs and chromatins were selected to serve as another control group. The perfect-matched sequence pairs form the positives for stable RNA-DNA hybrids. Purple dotted lines define the stable RNA-DNA hybrids with 95% confidence interval. (B) The proportions of RNA-DNA duplex-mediated LINE-1/ERV RNA-chromatin interactions in E14, H9 and HEK293 cells. The transposon RNA-chromatin interaction is duplex-mediated if a stable RNA-DNA hybrid duplex is formed in the paired sequence contexts. (C) The enriched overlap between R-loop loci and the duplex-mediated LINE-1 RNA-interacting DNA regions in hESCs. **** represents p <= 0.0001 in T-test showing the significance of the overlap by comparing with random sampling. (D) The consistence of RNA-DNA duplex-mediated LINE-1/ERV RNA-DNA interactions in H9 cells and differentiated HEK293 cells. (E) Two specific examples illustrating the loss of transposon RNA-chromatin interaction due to the genomic variations on the RNA-DNA hybrid sequences in HEK293 cells. An 18bp deletion and multiple single nucleotide variations respectively occur on the RNA-DNA hybrids in HEK293 for the two RNA-chromatin interactions. Bar plots represent the intensities of RNA-chromatin interactions (upper) and the minimal free energy (MFE) defining the binding strength of RNA-DNA hybrids (lower) in H9 and HEK293 cells for the two examples.

In principle, the existence of a stable RNA-DNA hybrid duplex would enable the de helix of double stranded DNA, accompanied by the formation of single stranded DNA fragment and an R-loop nucleic acid structure. DNA:RNA immunoprecipitation high-throughput DNA sequencing (DRIP-seq) data (Yan et al. 2020) in hESCs support the prediction of the formation of the RNA-DNA hybrid duplexes in transposon RNA-chromatin interactions (Figure 2C). R-loop signals are detectable in 68.3% of LINE-1 RNA interacting regions with stable RNA-DNA hybrid duplexes, statistically significant with P-value <0.0001 as compared with the negative control group. Comparisons between different human cell types demonstrate that about 70% hybrid-mediated LINE-1 or ERV RNA-chromatin interactions are consistent in H9 and differentiated HEK293 cells, free from cell identity (Figure 2D). Genomic variations (Lin et al. 2014) on the RNA-DNA hybrid duplexes abolish about 5% transposon RNA-chromatin interacting activity between H9 and HEK293 cells, as described in the following two specific examples (Figure 2D-2E). A stable RNA-DNA hybrid duplex with minimal free energy of −60.5 kcal/mol is formed that mediates the interaction between a LINE-1 RNA and a piece of DNA sequence on chromosome 19q13 in H9 cells. An 18bp deletion in the DNA sequence of this RNA-DNA hybrid in HEK293 cells leads to the reduction of the RNA-DNA binding strength with minimal free energy of −38.2 kcal/mol (Figure 2E). Meanwhile, in HEK293 cells the LINE-1 RNA-chromatin interaction is no longer detected at this region with the MARGI experiment data, although there are no differences between H9 and HEK293 cells in the expressions of other potential trans- or cis-regulatory factors around this region. Similarly, another case, where multiple single nucleotide variations occur in the ERV RNA and its targeting *LUADT1* gene body on the chromosome chr6q24 in HEK293 compared with H9 cells, demonstrates the loss of ERV RNA interaction is result from the weakening of the RNA-DNA hybrid (Figure 2E). All the above evidence reveals that the RNA-DNA hybrid binding mode is required for many transposon RNA-chromatin interactions.

### LINE-1 as an RNA scaffold inhibiting the expression of developmental genes in ESCs

Transposon RNAs of LINE-1, LTR, SINE and rRNAs target tens of thousands DNA loci in ESCs and HEK293 cells. These DNA loci distribute on gene promoter (−5kb~+1kb), intron and intergenic regions, as categorized in Figure 3A. Specially we studied transcriptional activation of the LINE-1 RNAs that target the gene promoter regions in mESCs. In mESC LINE-1 knockdown (LINE-1-KD) RNA-seq experiment, there are 388 up-regulated genes and few down-regulated genes after LINE-1 knockdown (Percharde et al. 2018). 73.45% (285) of the 388 up-regulated genes, named as LINE-1 trans-regulating genes, have LINE-1 targeting loci in their promoter regions (Figure 3B, Supplementary table S1). The LINE-1 targeting loci are enriched in the LINE-1 trans-regulating genes. These data support that LINE-1 RNAs play an important trans-regulating role of inhibition of gene expression in mESCs via the distal interactions with chromatins. The LINE-1 trans-regulating genes are significantly involved in the biological functions relative with development and cell proliferation regulation (Figure 3C). The functions are consistent with the cellular phenotypes of decreased proliferations, shortened cell-cycle S phase, and increased mortality in LINE-1-KD mESCs.

**Figure 3.**
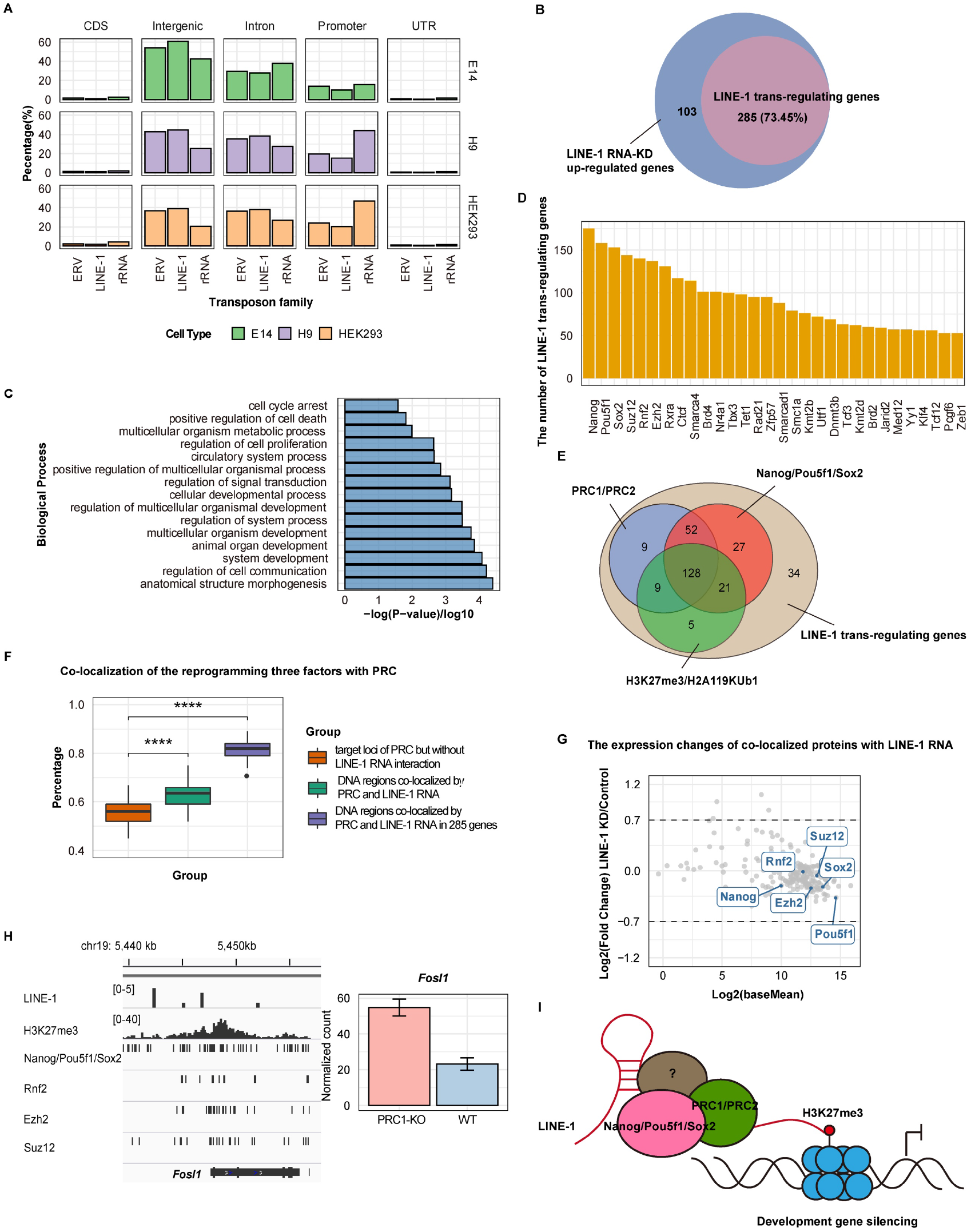
LINE-1 as an RNA scaffold inhibiting the expression of developmental genes in mESCs. (A) The genomic distribution of DNA loci targeted by LINE-1 RNAs, ERV RNAs and rRNAs. The gene promoter regions were extracted from −5kb to +1kb to transcription start sites of genes annotated in Ensembl. (B) The enriched proportion of LINE-1 trans-regulating genes. The LINE-1 trans-regulating genes are LINE-1-KD up-regulated genes with LINE-1 RNA targeting loci in their promoter regions. (C) GO enrichment analysis on biological process for LINE-1 trans-regulating genes. (D) The top 30 significantly co-localized proteins with LINE-1 RNAs in the promoter regions of the LINE-1 trans-regulating genes. The y-axis shows the number of LINE-1 trans-regulating genes co-bound by each protein. The binding sites of proteins were annotated by ChIP-seq data in GTRD (http://gtrd.biouml.org). (E) The venn plot for the co-localization of Polycomb repressive complexes (PRC) PRC1 (Rnf2) or PRC2 (Ezh2 and Suz12) with reprogramming three factors (Nanog, Pou5f1 and Sox2), and histone modifications H3K27me3 or H2A119KUb1 in the promoter regions of LINE-1 trans-regulating genes. (F) Significance analysis on the co-localization of the reprogramming three factors with PRC on LINE-1 trans-regulating genes. We randomly sampled 500bp-length DNA regions with targeting loci of PRC but without LINE-1 RNA interaction as the first control group. 500bp-length DNA regions co-localized by PRC and LINE-1 RNA were selected to be the second control group. The percentage bars of samples co-localized by PRC and the reprogramming three factors in the promoter regions of LINE-1 trans-regulating genes, and in the DNA regions of the two controls was plotted, respectively. **** represents the significance test with p-value <= 0.0001. (G) The expression changes of all the co-localized proteins with LINE-1 RNAs in the promoter regions of the LINE-1 trans-regulating genes after LINE-1 knockdown. No expression changes are found for the highlighted Polycomb group proteins and the reprogramming three factors. (H) An example of LINE-1-KD up-regulated gene *Fosl1* in the promoter region of which there are multiple loci co-bound by LINE-1 RNA, PRC and the reprogramming three factors. Integrative Genomics Viewer shows the ChIP-seq peaks of histone modification, PRC and the reprogramming three factors in GTRD and the interacting signals of LINE-1 RNA for this genes. Bar plot shows the significantly up-regulated expression of *Fosl1* after PRC1 knockout as well as LINE-1 knockdown in E14. (I) A gene silencing model that LINE-1 RNAs as well as Nanog/Sox2/Pou5f1 facilitate PRC-mediated repression of developmental genes in ESCs.

Using the most comprehensive protein ChIP-seq data in GTRD (Yevshin et al. 2017) (http://gtrd.biouml.org/), we analyzed the protein binding in the promoter regions of the LINE-1 trans-regulating genes. The promoter regions of these genes enrich the binding sites of many proteins including the reprogramming three factors (Nanog, Pou5f1, and Sox2) and Polycomb group proteins (Figure 3D). Polycomb repressive complexes PRC1 (Ring1, and Rnf2) and PRC2 (Eed, Ezh2, and Suz12) are one of the most important gene activation switches, which control cell fate by repressing gene activity. Previously studies reported that Oct3/4 and Sox2 mediate local engagement of PRC in gene promoters since PRC cannot directly bind to DNA (Endoh et al. 2008; Yu et al. 2012; Guo et al. 2018). The ChIP-seq data reveal that Polycomb group proteins bind to the promoter regions of 198 LINE-1 trans-regulating genes, 90.9% (180/198) of which are simultaneously co-occupied by the reprogramming three factors Nanog, Pou5f1, and Sox2 (Figure 3E, Supplementary table S1). LINE-1 RNAs significantly facilitate the co-localization of PRC and the three factors on the promoter regions of LINE-1 trans-regulating genes (Figure 3F). There are few expression changes for the reprogramming three factors and Polycomb group proteins when the LINE-1 trans-regulating genes are up-regulated after LINE-1 knockdown (Figure 3G). Therefore, LINE-1, in addition to the reprogramming three factors, mediates the PRC recruitment for the inhibition of hundreds of genes functionally related to development and cell proliferation. For example, *Fosl1* gene, which activates cell fate conversion from ESC to trophoblast, is co-occupied and co-repressed by LINE-1 RNA, the reprogramming three factors and Polycomb group proteins (Figure 3H) (Lee et al. 2018). Among the total 285 LINE-1 trans-regulating genes in mESCs, 229 (93.09%) of the 246 homologous genes in hESCs (H9) have LINE-1 RNA interacting loci in their promoter regions (Supplementary Figure S2A). In these 229 genes, 69 (82.14%) of the 84 genes targeted by Polycomb group proteins are simultaneously occupied by the reprogramming three factors in the promoter regions (Supplementary Figure S2B). The genes co-regulated by LINE-1 RNA, the reprogramming three factors, and the Polycomb group proteins in hESCs are related to biological functions of development and cell proliferation (Supplementary Figure S2C). The predicted minimal free energy of the interactive sequence contexts indicates that in both mESCs and hESCs the trans-regulation of LINE-1 RNAs, coordinating with the reprogramming three factors and the Polycomb group proteins, is not in the mode of RNA-DNA hybrid duplexes. In conclusion, we are proprosing a gene silencing model that LINE-1 RNAs, as well as Nanog/Sox2/Pou5f1, involve in PRC-mediated repression of developmental genes in ESCs (Figure 3I). In this way, LINE-1 functionally links to the core transcriptional regulatory circuitry for the stabilization of ESC identity.

### RNA-DNA hybrids guiding LINE-1 RNA-KAP1 coregulation on the transcription of LINE-1 lineages and *Dux* gene

The MARGI data and the comprehensive ChIP-seq data in GTRD support the co-localization of LINE-1 RNA and some proteins in the promoter or gene body regions, where the LINE-1 RNAs interact with chromatins via the mode of RNA-DNA hybrid duplexes (Figure 4A). These co-binding proteins include Kap1 as well as Zeb1, Smarcad1, Gli2, Tet1, Dppa2, and ZNF486. Previous studies observed that in ESCs, LINE1 acts as a nuclear RNA scaffold recruiting Kap1 to repress *Dux*, an activator of the transcriptional program in zygotic genome activation (De Iaco et al. 2017; Percharde et al. 2018). Our finding explains how LINE-1 RNA-Kap1 targets *Dux*. The R-loop signals (Sanz et al. 2016) and the minimal free energy (MFE) of RNA-DNA hybrid support it is the RNA-DNA hybrid duplex in the LINE-1 RNA and chromatin interactions that guides the LINE-1 RNA-Kap1 complex for the recognition on *Dux* gene (Figure 4B). The protein binding and histone modification data and the PRC-KO experiment data further indicate that PRC acts as an essential partner for the binding and repression on the *Dux* gene that is co-regulated by the LINE-1 RNA and Kap1 in mESCs (Figure 4B, Figure 4C). Besides *Dux*, the RNA-DNA hybrid duplex mediates the LINE-1 RNA-Kap1 for the binding and inhibition on some other genes relative with ESC differentiation (Figure 4C). Among these genes, *Adcy2* plays an important role in embryonic development including cell migration, proliferation, and differentiation (McCallie et al. 2019). And *C130026I21Rik*, a homolog for the gene *SP140* in human, participates in biological process of cell-cell adhesion that is crucial to maintain tissue morphogenesis and regulate cell migration and differentiation during development (Kashef and Franz 2015; Karaky et al. 2018).

**Figure 4.**
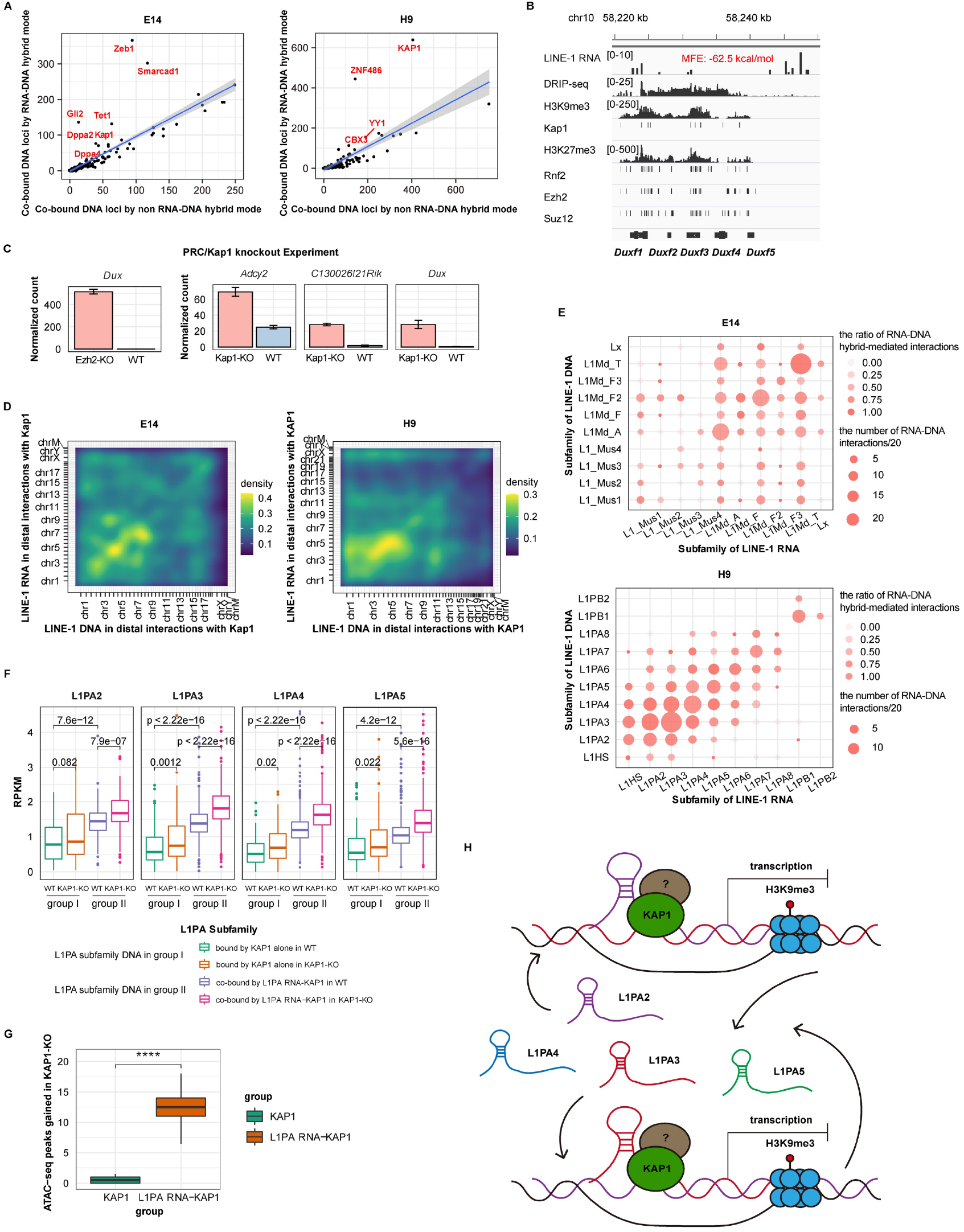
RNA-DNA hybrids guiding LINE-1 RNA-KAP1 coregulation on the transcription of LINE-1 lineages and *Dux* gene in ESCs. (A) The LINE-1 RNA-protein co-binding DNA loci by RNA-DNA hybrid mode vs. non RNA-DNA hybrid mode in E14 and H9 cells. X- and y-axis show the DNA loci numbers. Each scatter represents a co-binding protein with LINE-1 RNAs. Red color highlights the proteins that prefer to co-localize in the DNA loci with RNA-DNA hybrid mediated LINE-1 RNA. (B) The co-localizing signals on the *Dux* family members. Integrative Genomics Viewer presents the binding signals of PRC and Kap1, the H3K27me3/H3K9me3 histone modification peaks, the LINE-1 RNA interacting signals, DRIP-seq R-loop signals, and the minimal free energy (MFE) for RNA-DNA hybrids on the *Dux* family members. (C) The significant rescue of repression for *Dux*, *Adcy2* and *C130026I21Rik* expression in PRC1/Ezh2 or Kap1 knockout experiments in E14. LINE-1 RNA and Kap1 protein co-occupy the promoter or gene body regions of the three LINE-1 trans-regulating genes. (D) The chromosomal coordinate heatmaps of distal interactions between LINE-1 RNA and LINE-1 DNA with a co-binding partner of KAP1 in E14 and H9 cells. The whole genome is in 10Mb/bin resolution. (E) The numbers of distal interactions between LINE-1 RNA and LINE-1 DNA with a co-binding partner of KAP1 in LINE-1 subfamily level. The size of the nodes presents the interaction numbers between different LINE-1 subfamilies. And the color of the nodes presents the ratios of RNA-DNA hybrid-mediated interactions. (F) LINE-1 RNA-KAP1 coregulation on the transcription of L1PA subfamilies in H9 cells. We divided the each L1PA subfamily DNA into the group II that are co-bound by L1PA RNA-KAP1 via the RNA-DNA hybrids and the group I that are bound by KAP1 alone. The boxplots show the expression of the two groups of L1PA subfamilies in H9 cells and the sequent changes in KAP1 knockout. The L1PA subfamilies that are co-bound by L1PA RNA-KAP1 in the group II are significantly over-expressed after KAP1 knockout. And the expressions of the group II are significantly higher than that of group I before KAP1 knockout. P values are calculated based on Wilcox test in default parameters. (G) The changes of chromatin accessibility of the L1PA DNA regions that are co-bound by L1PA RNA-KAP1 and the L1PA DNA regions that are bound by KAP1 alone. The boxplots show the changes of chromatin accessibility after KAP1 knockout in H9 cells. P value is calculated based on T-test in default parameters. (H) The RNA-DNA hybrid guiding LINE-1 RNA-KAP1 coregulation model for LINE-1 subfamily transcription stabilization via a negative feedback loop.

The high-throughput MARGI data also reveal many distal interactions between LINE-1 RNA and LINE-1 DNA with a co-binding partner of KAP1 (Figure 4D). The interactive LINE-1 RNAs and LINE-1 DNAs are dominantly derived from subfamilies of L1PA2, L1PA3 and L1PA4 in human and L1Md-A, LMd-F2, and L1Md-T in mouse. Cross interactions occur within subfamily members and among different subfamily members (Figure 4E). The RNA-DNA hybrid duplex mediates 72.1% (1,263/1,752) and 81.2% (1,508/1,858) of such LINE-1 RNA-KAP1 co-binding events to the LINE-1 subfamily DNAs in hESCs and mESCs, respectively (Figure 4E). A feedback structure is physically formed in the transcription program of these LINE-1 DNAs with two key feedback components of the interactive LINE-1 subfamily RNAs and the co-binding KAP1. The LINE-1 DNAs of L1PA subfamilies that are co-bound by LINE-1 RNA-KAP1 via the RNA-DNA hybrids are significantly over-expressed after KAP1 knockout in hESCs. In contrast, there is no significant change in the expression of the control L1PA subfamily members that are bound by KAP1 alone (Figure 4F). The result demonstrates that RNA-DNA hybrid guides L1PA RNAs to function in the LINE-1 RNA-KAP1 complex for trans-repression of the transcription of L1PA lineage in hESCs. The chromatin accessibility of the L1PA DNA regions co-bound by LINE-1 RNA-KAP1 significantly increases after KAP1 knockout in hESCs, while there is no change in the chromatin accessibility of the control L1PA subfamilies that are bound by KAP1 alone (Figure 4G). The result indicates that interactive L1PA RNAs play a potential function related to chromatin opening of L1PA DNAs. Such a pattern of chromatin accessibility that impact protein binding to L1PA subfamily DNAs is consistent with the tendency of L1PA transcriptional changes in Figure 4F. The above evidence supports the active functions of both the LINE-1 subfamily RNAs and the co-binding KAP1 in the feedback structure of L1PA transcription program. The trans-regulation of LINE-1 RNA-KAP1 activates a negative feedback loop stabilizing the transcription of LINE-1 subfamilies (Figure 4H). Therefore, the L1PA RNA-KAP1 presents weaker inhibition on L1PA expression than KAP1 alone (Figure 4F).

## Discussion

Recent studies have highlighted the benefits of transposon RNAs to 3D genome structure, chromatin accessibility and heterochromatin formation (Jachowicz et al. 2017; Hao et al. 2020; Liu et al. 2020; Lu et al. 2021). In particular, the advanced high-throughput experimental biotechnologies exploring RNA-chromatin interactions are enabling the deep understanding of the potential function of transposon RNAs on DNA regulation in cells. When the ultra-short sequencing reads of about 20bp in GRID-seq (Li et al. 2017) limit the accurate recognition of complex repeatitive elements, the MARGI technology (Sridhar et al. 2017) with 150bp-length sequencing reads is conducive to the profiling of transposon RNAs and their distal targeting DNA loci. We performed MARGI experiment in mESCs and utilized in-house and public high-throughput data for analysis on trans-regulation of transposon RNAs. As conclusions, the transposon RNAs generally interact with chromosomal DNA in coding and intergenic regions via the RNA-DNA hybrid duplex mode or the mode of RNA scaffold recruiting DNA-binding proteins. It is worth noting that we improve some original ESC potency regulation models during mammalian early development. For examples, LINE-1 RNA as an essential RNA scaffold is involved in the co-regulation model of the three reprogramming factors and Polycomb repressive complexes. And in the way of RNA-DNA hybrids, LINE-1 acts as a guide RNA to mediate KAP1 or KAP1-PRC for expression inhibition of LINE-1 and some important developmental genes. The updated new models support that the transposon RNAs participate in the trans-regulation in the core transcriptional regulatory circuitry for the stabilization of ES cell identity.

In addition to the function of trans-regulating gene inhibition, we find transposon RNAs are required for many cell functions through RNA-chromatin interactions. For example, the binding sites of proteins CTCF and Smarcad1 are enriched in the LINE-1-interacting DNA regions. Previous studies revealed CTCF had a key role in the establishment of 3D chromatin structure during transcription and embryogenesis (Tang et al. 2015; Chen et al. 2019) and RNAs stabilized the binding between CTCF and chromatin (Saldana-Meyer et al. 2019). Our evidence that LINE-1 RNAs and CTCF protein co-localize on the chromosomal regions also indicates the involvement of the transposon RNAs in the establishment of 3D chromatin structure (Lu et al. 2021). DNA helicase Smarcad1 possesses intrinsic ATP-dependent nucleosome remodeling activity. The co-localization of LINE-1 RNA and Smarcad1 in the heterochromatin regions is Kap1-dependent and enriches with H3K9me3 loci in mESCs. We suggest that LINE-1 RNAs may be involved in the heterochromatin organization in the form of Smarcad1-LINE1-Kap1 in mouse, similar to the function of ERV RNAs (Hao et al. 2020; Liu et al. 2021).

Although the majority of chromatin-interacting repetitive elements are transposon RNAs, ribosomal ribonucleic acids (rRNAs), composed largely of short tandem repeats, also exhibit distal interactions with chromatins. We observe that there is a preference for low complexity regions with simple G-rich or (CGG)n repeat units in the interactive DNA regions of rRNAs. As a characteristic of G-quadruplex, G-rich content contributes to genomic instability during DNA replication (Garcia-Muse and Aguilera 2016). The (CGG)n repeat, being the underlying DNA structure of rare fragile sites, also promotes the breakage of DNA double strands (Durkin and Glover 2007). Secondly, up to 85% of the rRNA-interacting regions contain known chromosome fragile sites (Kumar et al. 2019) and there is a tendency of co-localization of rRNA and homologous recombination repair protein BRCA1 in the chromatin regions. The evidence indicates the potential relationship between rRNA-chromatin interactions and genomic instability.

## Methods

### Mouse embryonic fibroblasts cell culture

E13.5 primary MEFs derived from pooled CD1 embryos (mixed sex) were cultured at 37℃, 5% CO# in MEF medium (high glucose DMEM with HEPES (Thermo Fisher Scientific, #12430054), 10% FBS (Gibco, #10270-106), 2mM GlutaMAX supplement (Gibco, #35050061) and 2mM 100 U/mL penicillin-streptomycin (Gibco, #15140122)), and used within 4 passages of initial derivation. MEFs were treated with mitomycin C (StemCell Technologies, #73274) for 2.5-3 hours before they were used as the feeder layer.

### Mouse embryonic stem cell culture

Mouse E14Tg2A (E14) ES cells (male) were obtained from ATCC (#CRL-1821) and used for mESC pxMARGI experiments. ESCs were cultured at 37℃, 5% CO# on 0.1% gelatin (Millipore, #ES-006-B)-coated plates in ES-FBS and MEF (treatment with mitomycin C)-conditioned culture medium (high glucose DMEM with HEPES (Thermo Fisher Scientific, #12430054), 15% FBS (VISTECH, #SE200-ES), 0.1mM non-essential amino acids (Gibco, #11140050), 2mM GlutaMAX supplement (Gibco, #35050061), 1 × Nucleosides (Millipore, #ES-008-D), 2mM 100 U/mL penicillin-streptomycin (Gibco, #15140122), 0.1mM 2-Mercaptoethanol (Millipore, #21985023) and 1,000 U/ml LIF supplement (Millipore, #ESG1107)). Sensitive alkaline phosphatase (AP) activity testing of E14 ES cells revealed a highly pure population of mESCs. To remove MEF cells, we firstly dissociated cells with Trysin (Gibco, # 25300054). Then, we distributed each plate E14 ESCs suspension onto two new gelatin-free plates with enough media to cover the bottom for 1 hour and separated from the MEF feeder layer by differential sedimentation of the MEF and E14 ESCs.

### MARGI experiment

The MARGI experiment in E14 ESCs was performed as previously described (Sridhar et al. 2017). Briefly, the 4-5 million cells were crosslinked by 1% formaldehyde (Thermo, #28906). Chromatin was fragment using HaeIII (NEB, #R0108M) and subsequently stabilized on streptavidin T1 beads (Thermo, #65601) after cell lysis and biotinylation of proteins (Thermo, #21334). Then, RNA and DNA ends were prepared for ligation. First, a non-templated dA tail was added using Exo(−) Klenow (NEB, #M0212S) to 3’-P presenting in the RNA that was dephosphorylated using T4 PNK (NEB, #M0201S). Linker and RNA ligation was performed by ligating pre-adenylated linker to the 3’-OH of RNA using RNA ligase2, truncated KQ (NEB, #M0373L). Reaction was carried out at 22℃ for 6 hours followed by 16℃ overnight. Second, DNA ends were phosphorylated using T4 PNK (NEB, #M0201S) and T4 DNA ligase (NEB, #M0202M) were used for proximity ligation by 16℃ overnight. After reverse crosslinking and DNA/RNA extraction, the RNA was reverse transcribed using Supercript IV (Invitrogen, #18090050) and purified the single strand DNA without biotin using Silane beads (Invitrogen, #37002D). Then, purifying DNA was circularized and digested by restriction enzyme BamHI (NEB, #R3136S). Finally, the right and left flanking sequences of the linker were split as primers for amplification and sequencing.

### Detection for distal transposon RNA-chromatin interactions from MARGI data

1. Alignment of DNA and RNA reads The two ends of RNA-DNA paired reads were separately processed. Firstly, reads processing, including removing adaptor, filtering low-quality and duplication reads, was performed according to previously published pipeline (Sridhar et al. 2017). Then, we ran STAR (Dobin et al. 2013) using the following options: --runThreadN --genomeDir --alignEndsType Extend5pOfRead1 --seedSearchStartLmax 25 --outFilterScoreMin 21 --outFilterScoreMinOverLread 0.13 --outFilterMatchNmin 21 -- outFilterMatchNminOverLread 0.13 --alignIntronMax 1 and BWA (Li and Durbin 2009) MEM using the following options: −k 15 –T 15 for DNA and RNA ends, respectively.
2. The RNA ends of RNA-DNA paired reads were mapped to transposon element regions that were annotated in UCSC RepeatMasker.
3. Mappable DNA reads with multiple hits were removed.
4. We filtered the mappable DNA reads that were annotated in heterochromatin regions via H3K9me3 and HP1a ChIP-seq data (GSE57092, GSE39579, ENCODE). Exceptionally the LINE-1 DNAs in heterochromatin regions co-bound by LINE-1 RNA-KAP1 were retained.
5. The definition of the term “distal” followed the rule that the chromosomal coordinates between the mappable RNA end and DNA end of RNA-DNA paired reads should be more than 5kb apart.
6. The DNA loci targeted by transposon RNAs, spanning ~200bp region, are defined by the clusters of distal RNA-DNA paired reads.

### RNA-DNA hybrid prediction between the sequence contexts of a transposon RNA-chromatin interaction pair

10,000 DNA sequences outside the transposon RNA targeting regions were randomly sampled for 100 times to compose the pseudo transposon RNA-DNA pairs as control. And 20,000 confident distal interaction pairs between non-transposon RNAs and chromatins were selected to serve as another control group. The repeat information was annotated by UCSC RepeatMasker. The perfect-matched sequence pairs form the positives. Predictions of triple helix and DNA/RNA hybrid duplex structure were performed using Triplexator and RNAhybrid (Kruger and Rehmsmeier 2006; Buske et al. 2012) on the above datasets. A 50bp sliding window along transposon RNA was utilized for RNAhybrid prediction. The minimal free energy defines the binding strength of RNA-DNA hybrid duplex.

### Peak calling from R-loop and histone modification/protein ChIP-seq data

We re-analyzed R-loop data were downloaded from NCBI (GSE145964, GSE70189). The raw data processing was the same as pxMARGI data and uniquely aligned to hg19 or mm10 genome by STAR (Dobin et al. 2013) with default parameters. MACS2 (Zhang et al. 2008) with the --broad parameter was used for peak calling.

H3K9me3 stable peaks in H9 (ENCODE) and H2A119KUb1 peaks in mESC (GSE145964) were used, directly. H3K27me3 (ENCODE, Epigenome), H3K9me3 (ENCODE, GSE57092) and HP1a (GSE57092, GSE39579) ChIP-seq data of the above mentioned were downloaded. For raw FASTQ data, we used Bowtie2 (Langmead and Salzberg 2012) or BWA (Li and Durbin 2009) with the default parameters for alignment of ChIP-seq data. MACS2 (Zhang et al. 2008) was used for peak calling from BAM or BED files.

### RNA-seq data processing

Public available FASTQ files from H9, HEK293 and E14 RNA-seq experiments from the Gene Expression Omnibus (GSE103715, GSE43572, GSE54106) were removed PCR duplicated reads and ran BWA (Li and Durbin 2009) MEM using the same options as the RNA ends of read pairs from MARGI experiments to reduce technical bias. For Kap1-KO RNA-seq data (GSE99215) in H9, reads processing was performed using the same options and mapped to mm10 or hg19 genome using STAR (Dobin et al. 2013) with the following parameters: --outFilterMultimapNmax 100 -- outFilterMismatchNoverReadLmax 0.04 --alignIntronMax 1000000 -- alignMatesGapMax 1000000, that enabled to output multiple alignments for a read. Normalized reads count per gene in Kap1-KO experiment were calculated based on the total mappable reads of Kap1-KO and WT data. For PRC-KO RNA-seq data (GSE66814, GSE132753), the results of differential expression analysis were used, directly.

### Genomic annotations on aligned reads or called peaks

Genomic information, including promoter (+5/-1kb from TSS), CDS, UTR, intron and intergenic regions, and gene symbols, was extracted from Ensembl (Homo sapiens.GRCh37.75, Mus musculus.GRCm38.79). The aligned reads and called peaks from high-throughput sequencing data were annotated with the genomic information. The biomaRt package (Durinck et al. 2009) was used for the gene symbol conversion between human and mouse species.

### The definition of co-localization between two binding loci

The co-localization of binding sites of protein/histone modifications or R-loop regions was determined via BEDTools (Quinlan and Hall 2010) if these regions located within 500bp region of transposon RNA binding loci. The binding sites of transcription factors downloaded from GTRD (Gene Transcription Regulation Database) (Yevshin et al. 2017), that is the most complete collection of uniformly processed ChIP-seq data on identification of transcription factor binding sites for human and mouse.

### Gene Ontology and Pathway Enrichment Analysis

Gene Ontology enrichment analysis was conducted by David.

### Quantification and statistical analysis

All statistical analyses were performed in R/Bioconductor. All statistics were * p < 0.05, ** p < 0.01, *** p < 0.001, **** p < 0.0001. They were calculated by Wilcox test unless specially noted otherwise.

## Data access

The datasets supporting the current study have been deposited in the Genome Sequence Archive under the accession number CRA003989 linked to the project PRJCA004596.

## Acknowledgments

We thank professor Jie Ren and Yungui Yang for constructive criticism and fruitful discussion. We thank Dr. Minglei Shi, Dr. Yang Chen and Professor Michael Zhang for kind help on experiment protocol.

This work was supported by grants from the National Key R&D Program of China [2018YFC1003102 to C.Z. and 2018YFC0910402 to J.C.]; the Strategic Priority Research Program of the Chinese Academy of Sciences [E0XD842201 to J.C.]; the National Natural Science Foundation of China [32070795 to J.C. and 31900603 to L.R.]; the Youth Innovation Promotion Association of Chinese Academy of Sciences [2020104 to C.Z]; the Open Project of Key Laboratory of Genomic and Precision Medicine, Chinese Academy of Sciences.

## Author Contributions

J.C. and C.Z. envisioned the project. W.X implemented the experiment and performed the analysis. W.X. and J.C. wrote the paper. C.Z. and L.R. provided assistance in writing and analysis.

